# OdoBD: An online database for the dragonflies and damselflies of Bangladesh

**DOI:** 10.1101/804658

**Authors:** Md Nur Ahad Shah, Md Kawsar Khan

## Abstract

Combining scientific data over a long-time period is necessary to understand the diversity, population trends, and conservation importance of any taxa in a global and regional scale. Bangladesh is located in a biodiversity hotspot region, however, till date, only few animal groups has been extensively investigated at a nation-wide scale. Although being one of the earliest and well-known insect groups, the knowledge on Odonata of this region remains rudimentary and dispersed. To resolve this issue, we have developed an online database for the Odonata of Bangladesh. We have compiled data from our last four years field study, from previously published research articles, field guides, and also collected data from citizen scientists regarding Bangladeshi odonates. Odonata of Bangladesh database (http://www.odobd.org) contains information on morphology, abundance, gene and protein sequences, local and global distribution and conservation status of the Odonata of Bangladesh. The database also demonstrates gender specified photographs with descriptions for better understanding for the novice researchers and naturalists. Odonata of Bangladesh database provides a comprehensive source for meta-analyses in ecology, conservation biology, and genetic research.

## Introduction

The inscription of natural organisms can be regarded as one of the most valuable documents in the study of historical occurrence of organisms. Although museum records, which serve as a credible source of information about the diversity of biological organisms, keep voucher specimens; comparing and contrasting all these records from different geographical locations for large-scale analysis prove to be an exceedingly difficult task. To confront this challenge, different types of digital catalogs started to be developed since 1970. A number of consolidated online databases like VertNet (Guralnick & Constable 2010), iNaturalist (http://www.inaturalist.org/), IUCN Red List (http://www.iucnredlist.org/), and Atlas of Living Australia (https://www.ala.org.au/), with their geographical, physiological and biochemical information, are proving to be an essential source for large-scale analysis conducted by scientists and researchers from all around the globe (Pyke & Ehrlich 2010). The online databases focusing on invertebrates, especially insects are, however, lagging far behind (Schuh *et al.* 2010). Although, this number is gradually increasing by the day as more insect data are being digitized in online repositories like OdonataCentral (Abbott & Broglie 2005), AllOdonata (http://www.allodonata.com/), FreshWaterBiodiversity (http://data.freshwaterbiodiversity.eu/), and Global Biodiversity Information Facility (http://www.gbif.org/), in response to the growing needs worldwide. Along with these online databases containing information on species worldwide, regional databases like Butterflies of India (Kunte *et al.* 2018), Butterflies of Belgium (Maes *et al.* 2016) are currently emerging, providing more detailed insights on the extant species with their spatial and temporal information.

The order Odonata is one of the earliest and well-known insect groups, existing on all continents except Antarctica (Trueman 2007). These insects predominantly inhabit the tropical and subtropical climate zones (Dumont 1991). Adult Odonates are terrestrial in nature, found adjacent to water sources, whereas the immature stages are aquatic, inhabiting freshwater habitats of all kinds, ranging from permanent running waters like rivers and lakes to small temporary rain pools and puddles. Being a species specific to a certain type of habitat makes them an ideal candidate for monitoring the health of freshwater ecosystems. The taxonomic Order Odonata is divided into three suborders – Anisoptera, which encompasses dragonflies; Zygoptera, which includes damselflies; and Anisozypgoptera, which contains intermediary species between these two groups. Till now, A total of 6,265 species of Odonates under 600 genera have been reported globally (Schorr & Paulson 2018). A combined effort has been undertaken to digitize all the information available on the odonata worldwide, in order to make them readily available to the interested scientific community (Schorr & Paulson 2018). Additionally, there have been region-specific studies on the odonates in different parts of the world, specifically countries with a diverse range of Odonata (Joshi *et al.* 2018; Kipping *et al.* 2009).

Bangladesh is a small country with high Odonata diversity. Currently, more than a hundred species are known from Bangladesh. The largely unconsolidated information makes large-scale analysis and research involving Bangladeshi Odonates particularly challenging. Thus, we have developed an online database of all the known Odonates from different locations of Bangladesh to generate an integrated and widely accessible source to facilitate studies of ecology, conservation, and genetic analysis. Currently, we have amassed information of 103 different species from all over the country. The database, named Odonata of Bangladesh (http://www.odobd.org), contains information on morphology, habitat, abundance, gene and protein sequences, worldwide distribution and conservation status of the Bangladeshi Odonates and is updated on a regular basis. We have included gender specified photographs with descriptions for better understanding for the novice researchers and naturalists. This database will spread the knowledge of the Bangladeshi Odonates as well as will enhance the opportunities for ecological and genetic research on those species.

## Methods

### Geographic coverage

The database Odonata of Bangladesh assimilated geographical data of the Bangladeshi odonates from the whole country. For the development of the database, we have divided the country into seven major regions, namely Dhaka, Rangpur, Rajshahi, Sylhet, Chittagong, Barisal, and Khulna (Figure: 01). Our study was mainly focused on four specific regions serving as Odonata breeding hotspots – Dhaka, Sylhet, Chittagong and Khulna; which correspond to the central, north-eastern, south-eastern and south-western part of the country, respectively. These regions encompass nearly all of the distinct climates and waterbodies of the country. We did regular surveys throughout the year for the last four years (2012-2016) in those regions and updated the checklist, parts of which have been published previously (Khan 2015a, 2015b, 2017; Khan & Tuhin 2018). We incorporated our unpublished data and data from published research for those regions in the database as well. We collected data of the Odonata on the other parts of Bangladesh from our occasional surveys, from the data deposited by the citizen scientists in our database and from the previously published research articles.

**Figure 1:**
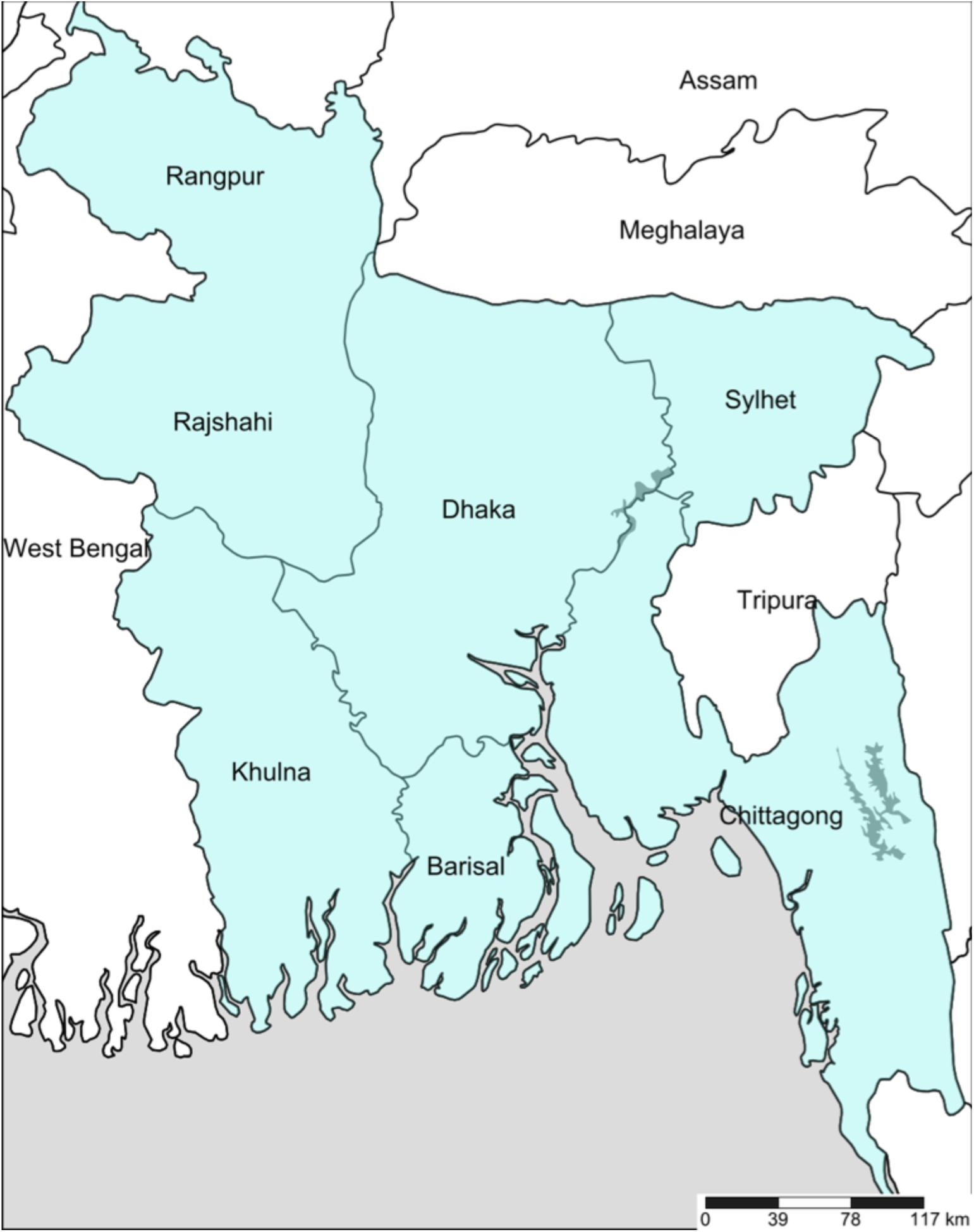
A reference map of different divisions of Bangladesh.

### Photographs

We captured most of the photos deposited on the database in the last four years (2012-2016). We captured various identification features of the dragonflies by a Canon 600D camera using a 55–250 mm lens. Most of the photos were taken in natural habitats from 0800-1700 hours (Khan 2015a). A few species were also photographed from previously collected specimens (Khan 2015a, 2015b). In addition to our photo database, we have also collected photo evidence from citizen scientists, providing their credit in the photos. We identified the photographs of the dragonflies using the identification keys and distinguishing features developed by the earlier entomologists (Asahina 1967; Fraser 1933, 1934, 1936; Lahiri 1987; Mitra 2002; Nair 2011; Subramanian 2009). We collected photographs of males and females and their different color morphs. We also took photographs of their life history such as emerging, perching, foraging, mating, oviposition, territorial fight etc.

### Data resources

Flight season, abundance and preferred eco-system were surveyed on these species to portray the regional diversity and prevalence. In all subsequent analyses, any species considered to be vagrant, with only one sighting was removed. Common name, IUCN status and global distribution data were collected from International Union for Conservation of Nature and Natural Resources database (http://www.iucnredlist.org/). In conjunction with the information, gender-specific morphological features like abdomen and wing size, distinct wing spots and colors were added from our literature review (Nair 2011; Subramanian & Gadgil 2009). For the purpose of genomic and proteomic studies, gene and protein sequences were also included in the database. As genomic and proteomic studies of odonates are understudied, the number of sequenced genes and proteins is very low. We included the most sequenced gene in the odonates cytochrome oxidase and its corresponding protein cytochrome c oxidase (EC 1.9.3.1) to study the phylogenic relationships. The gene and protein sequences for the species were collected from The National Center for Biotechnology Information database (http://www.ncbi.nlm.nih.gov/), UniProt database (http://www.uniprot.org/). Furthermore, we have included all the previous studies done on odonates in this region along with local field guides, in our bibliography section.

### Data structure

The OdoBD database contains records of species occurrences with their location and flight season. Taxonomic, ecological, and physiological information of Bangladeshi Odonata are included in the database. The gene and protein sequences, that are currently available, are also stored and provided in a separate tab. Thus, the following data are stored for each accession: Taxonomy, classification, general information (common name, scientific name, abundance, flight season, local and global distribution, and IUCN status, description, and gene and protein sequences. Photographs of male, female, foraging and reproductive behavior (copula, oviposition, tandem) are provided when available. A map of local distribution of each species is also included. General information on the difference between dragonfly and damselfly, their morphology, habitat, reproductive behavior, predator and prey interrelationship and conservation status are included under category ‘Biology’. The bibliography section was updated with a list of the previously published article on the Odonata fauna of Bangladesh. A common portal was created for citizen scientists to interact with the OdoBD database management team to submit sightings of the Odonata of Bangladesh.

### Database design

The MySQL database runs on a Percona server (webserver cpsrvd ver.11.x). Data are stored in the relational tables of a MySQL ver. 5.x database. A graphical user interface (GUI) named phpMyAdmin (v4.3.8) was installed in the server for managing MySQL tables and data. The GUI is accessible through all class of browsers regardless of operating system, though it has been most intensively tested using Mozilla Firefox and Google Chrome.

## Results

### Taxonomic Coverage

#### Kingdom

Animalia

### Class

Insecta

### Order

Odonata

### Suborders

Anisoptera; Zygoptera

### Families

Aeshnidae, Gomphidae, Libellulidae, Macromiidae; Calopterygidae, Chlorocyphidae, Coenagrionidae, Euphaeidae, Lestidae, Platycnemididae

### Genera

*Anaciaeschna, Anax, Gynacantha, Ictinogomphus, Macrogomphus, Megalogomphus, Orientogomphus, Paragomphus, Acisoma, Aethriamanta, Brachydiplax, Brachythemis, Bradinopyga, Camacinia, Cratilla, Crocothemis, Diplacodes, Hydrobasileus, Indothemis, Lathrecista, Lyriothemis, Macrodiplax, Neurothemis, Onychothemis, Orthetrum, Palpopleura, Pantala, Potamarcha, Rhodothemis, Rhyothemis, Tetrathemis, Tholymis, Tramea, Trithemis, Urothemis, Zyxomma, Epophthalmia; Matrona, Neurobasis, Vestalis, Aristocypha, Libellago, Aciagrion, Agriocnemis, Amphiallagma, Argiocnemis, Calicnemia, Ceriagrion, Ischnura, Mortonagrion, Paracercion, Pseudagrion, Dysphaea, Euphaea, Lestes, Coeliccia, Copera, Onychargia, Prodasineura*

### Database summary

There was a total number of 103 species recorded from all over the country during the study period. The number of recorded Anisoptera (Dragonfly) was 58; in which the family Libellulidae (the skimmers or perchers) were the greatest in number with 45 species on record and the family Aeshnidae (the hawkers) and Gomphidae (clubtail dragonflies) had 6 species each (Figure 2). *Epophthalmia vittata* was the only existing dragonfly in this region belonging to the family Macromiidae. The number of Zygoptera (Damselfly) was 45; where species of six different families were found (Figure 2). The family with the highest documented species was Coenagrionidae with 27 species and the family with least recorded species was Lestidae with only one species (*Lestes praemorsus*). Another low diverse recorded family was Chlorocyphidae with two species only. The family Calopterygidae and Euphaeidae had three recorded species each whereas nine species were recorded from Platycnemididae family. The sub-order Anisozypgoptera, which contains intermediary species between these two groups, has only two living species and none of them are found in Bangladesh.

**Figure 2:**
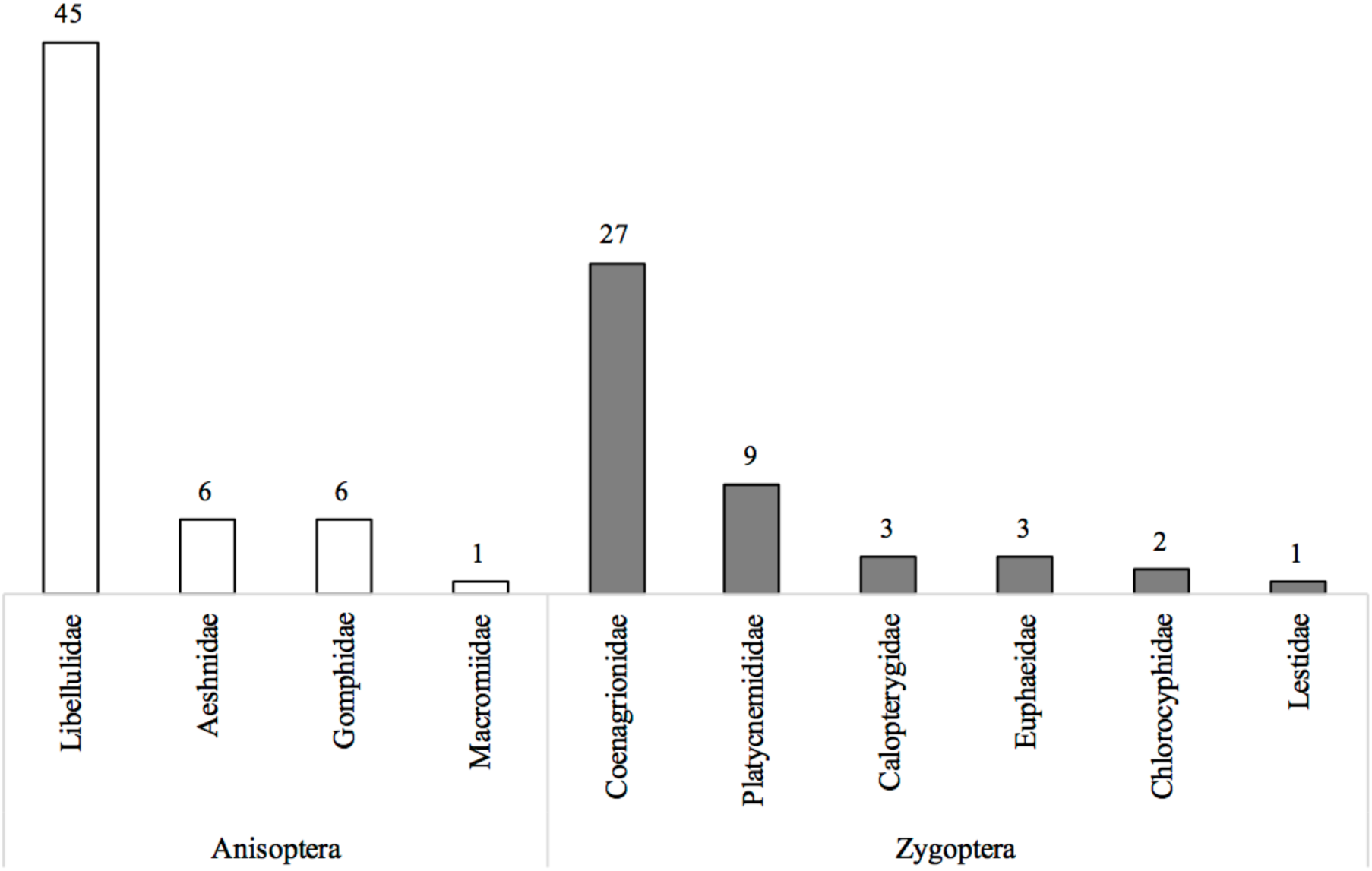
Number of extant species of different families of Odonates. White bars represent the sub-order Anisoptera and Grey bars represent the sub-order Zygoptera.

Currently, the database contains male specific photos of 83 species and 49 species’ female specific photos. Additional photos like perching, mating, oviposition are also included. Photos of rest of the species will gradually be added upon their availability.

### Temporal summary and conservation status

Among the seven divided divisions, Sylhet and Chittagong were found to be regions with the most species diversity. In Sylhet around 54 and in Chittagong around 45 different Anisoptera species were encountered (Figure 3). For Zygoptera, there were 34 encounters in Sylhet and 36 in Chittagong. Dhaka and Khulna have a moderate level of species diversity with a total sighting of 48 and 49 different odonate species respectively (Figure 3). The rest of the three regions have a lower number of species sightings. The species *Anax indicus* was only spotted in Rajshahi and Sylhet. *Macrodiplax cora* species was only seen in Chittagong and Khulna.

**Figure 3:**
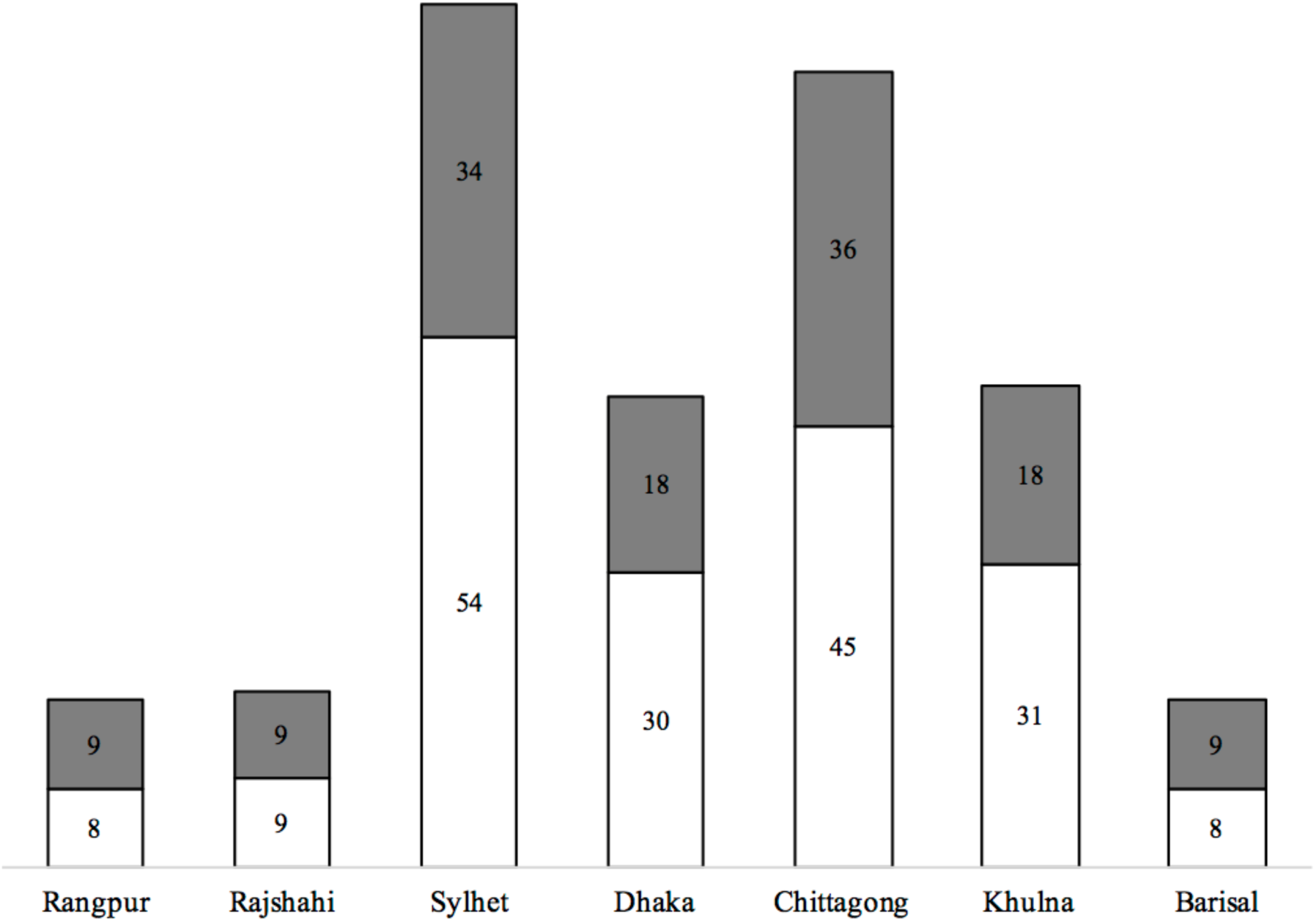
Number of extant species in different regions of Bangladesh. White bars represent the sub-order Anisoptera and Grey bars represent the sub-order Zygoptera.

The IUCN Red List status analysis showed, among the documented 101 species, 93 belongs to Least Concern (LC) category of which 51 species were Anisoptera whereas 42 were Zygoptera (Figure 4). Seven species were recorded under the category of Data Deficient (DD), in which all the species belonged to the sub-order Anisoptera (Aeshnidae 2 and Gomphidae 4). One documented species *Indothemis carnatica* belongs to Near Threatened (NT). The rest of the two Zygoptera species, namely *Matrona nigripectus* and *Agriocnemis Kalinga*, have not yet been assessed by IUCN.

**Figure 4:**
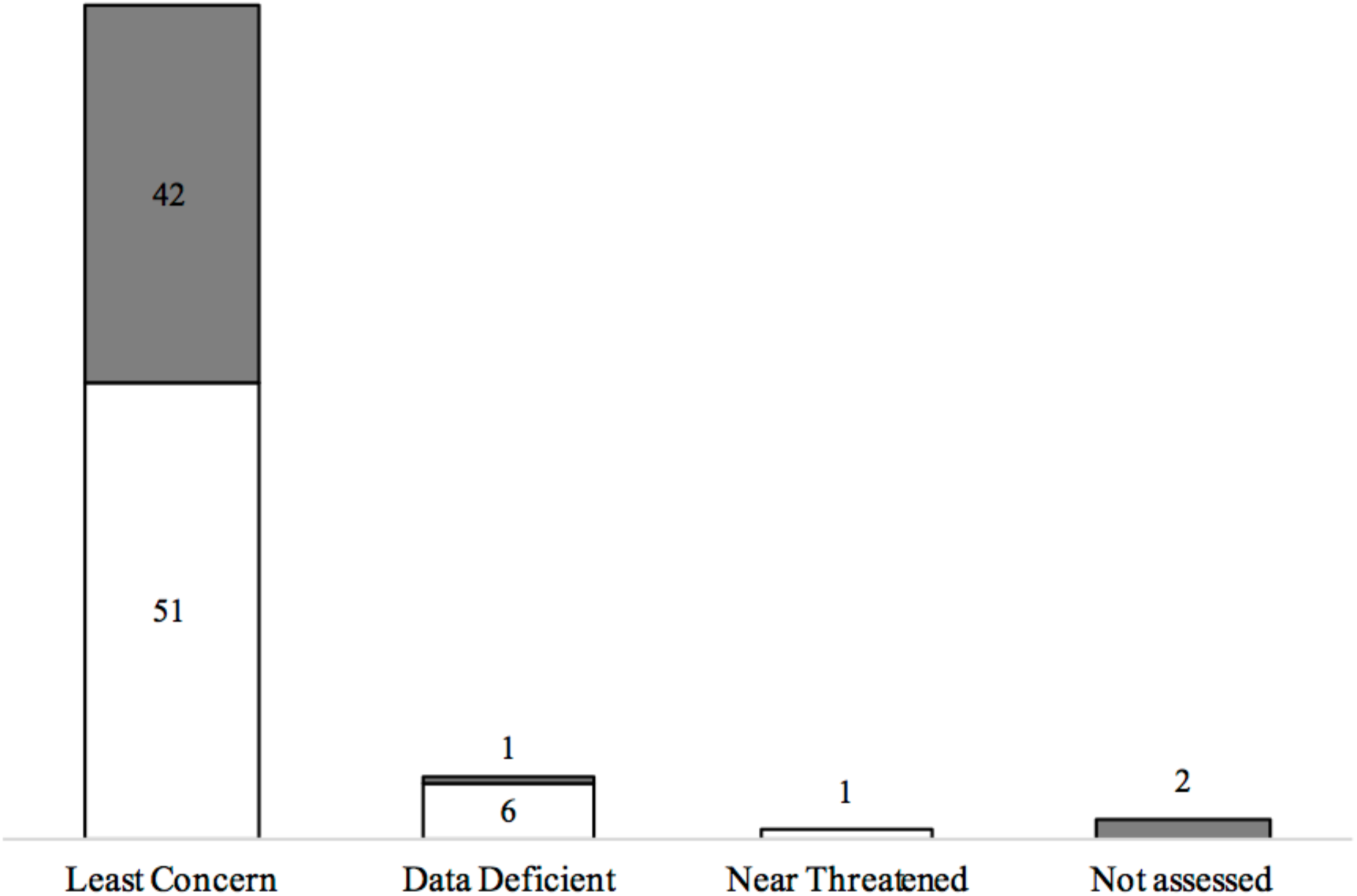
The IUCN status of different existing species in Bangladesh. White bars represent the sub-order Anisoptera and Grey bars represent the sub-order Zygoptera.

The frequency of sightings was observed for each species and is stated accordingly in the database. 51 species were found to be common in their respective zones, among *A. kalinga* is stated as not assessed by IUCN. Uncommon occurrence was noted for 25 species. Three species were locally common (*Cratilla lineata, Macrodiplax cora, Calicnemia imitans*). 19 species were found to be rarely occurring, in which four have Data Deficient and one species is nearly threatened. Three species were sighted in extremely rare cases where all three of them possess the Data Deficient status.

### Flight season and Genetic data

Among the 103 species, the flight season data was collected for 101 species. Due to the geographical location, Bangladesh has a temperate climate and has six seasons. Each season is comprised of two months, some of them are short while some flow into other seasons. The six seasons and their ranges are – Summer (April-May), Monsoon (June-July), Autumn (August-September), Late Autumn (October-November), Winter (December-January) and Spring (February-March). Eleven species were found to be abundant all year long. The sightings of odonates, peak with the maximum in the month of June, which is the starting of the season monsoon, and continues until October, that is the mid of late autumn (Figure 5). Their prevalence starts to decline with the least number of sightings during the winter season.

The genetic and proteomic data were collected from The National Center for Biotechnology Information database (http://www.ncbi.nlm.nih.gov/) and the UniProt database (http://www.uniprot.org/) for the gene and protein sequences. Sequences for the gene cytochrome oxidase and its corresponding protein cytochrome c oxidase (EC 1.9.3.1) for the available species (Currently 67 gene and protein sequences each) are continuously being incorporated into the database with their accession id as soon as they are made available on the different public databases.

**Figure 5:**
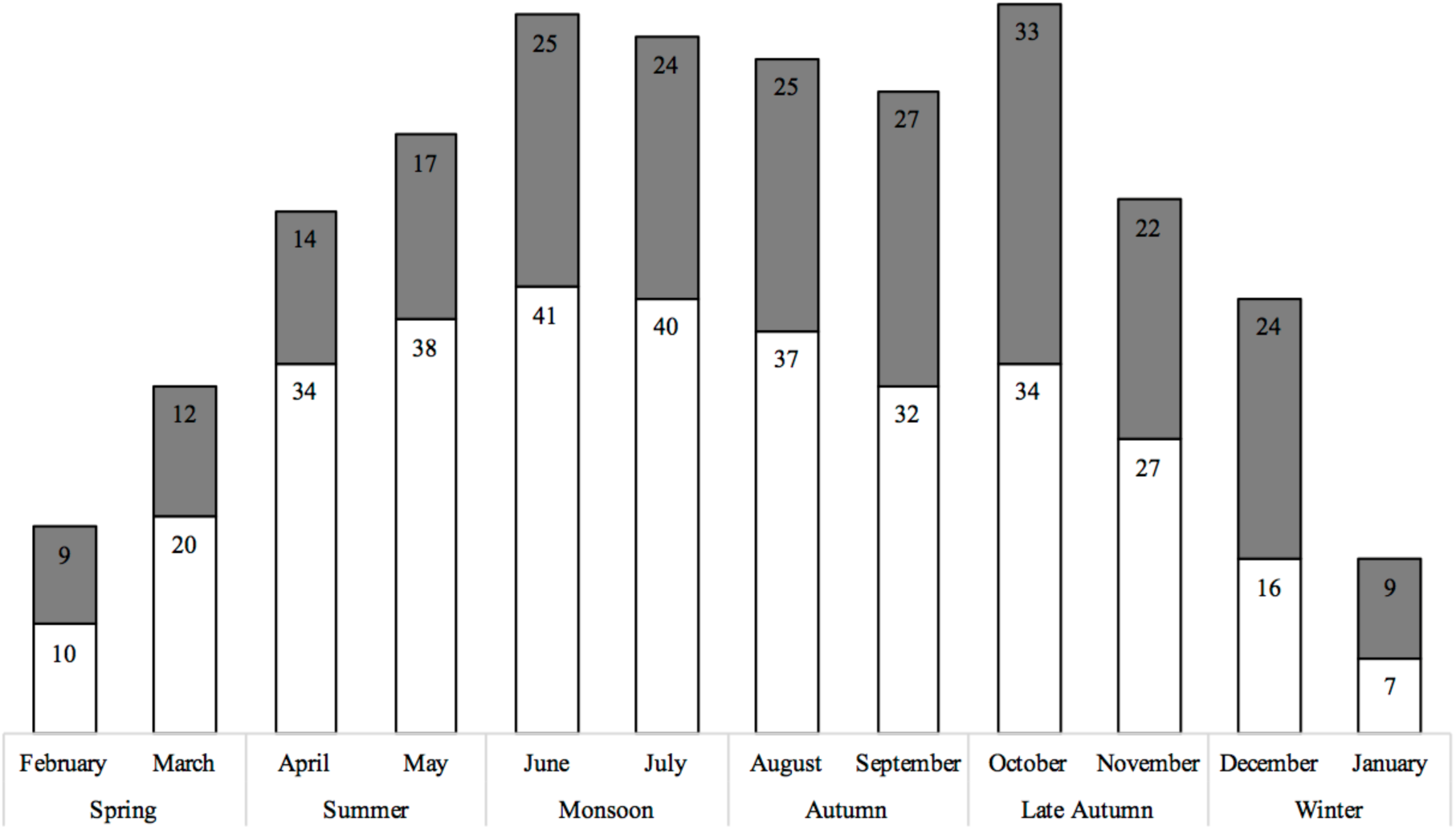
Number of species based on their flight pattern in different seasons of Bangladesh. White bars represent the sub-order Anisoptera and Grey bars represent the sub-order Zygoptera.

### Data accuracy and future updates

The data in OdoBD were entered by the first author (MNAS). Then entries were double-checked completely or spot-checked with a random sub-sample by MKK. Over 95% of such double-checked entries were correct. However, not all data entries were verified by a second person. Thus, some low level of data entry errors remains. Technical support and web development were done by MNAS and Borhan Uddin. Although the study time period mentioned was from 2012-2016, reports and sightings from 2017 onwards are still being recorded for this database and recent articles are being added whenever they are made available. We will continue to use feedback from peer reviewers and users of the database, to update and correct the database.

## Discussion

The storage and accession of the large amount of data for scientific research can be attained by the means of the universal electronic database. This database ‘OdoBD’ can be the source for various researchers for the faunistic, biogeographic and systemized research on the odonates of Bangladesh. It is also useful for spatial research on the odonata fauna as it encompasses numerous parameters like preferred eco-system, relative abundance, flight season, local and global distribution for each species. With the aid of different tools from the Geographical Information System (GIS), it is now possible to generate distribution maps of each species across a large area that can facilitate further research (Wadsworth & Treweek 1999). For Bangladesh, no such study along with data digitization has been carried out to date. We report this database as the first online and readily accessible checklist on odonata with additional information on each species from Bangladesh.

Our current compiled list of 103 species of odonata substantiates the rich faunal diversity of the country. Among the seven divisions, the most number of species was recorded in Sylhet and Chittagong, which contains most diverse water resources and also till date most of the study were concentrated in this region. Dhaka and Khulna have a moderate number of species diversity. The rest of the three regions – Rangpur, Rajshahi, and Barisal were documented with lowest species diversity. This low Odonata diversity can be attributed to the availability of the small amount of fresh water habitat and low data availability from these regions.

The Odonata database gathered data from grey literature, field guides, published articles and collected data from citizen scientists. There is a constant effort to keep the information updated along with their improved comprehensiveness, but biodiversity data are often quite sparse and can be geographically biased (Collen *et al.* 2008; Yesson *et al.* 2007). In cases, many of the data-collection sources under-represent certain areas that are in species-rich tropics. The lack of museum records on odonates in this region poses another big challenge in the acquisition of authentic documentations. Harnessing citizen science in these under-studied regions to monitor and document Odonata biodiversity can vastly improve our current knowledge on these species (Silvertown 2009). The rapid expansion of the Internet and advancement of mobile computing have accelerated the number of citizen science projects in recent years. This platform has the potential to facilitate the collection and circulation of taxonomic data covering a wide geographical area at minimal expenditure. Our database provides such a platform for the contributions from citizen scientists across the nation. The species submission portal allows the users to enter data with photographic evidence that are directly sent to the online server. Unusual sightings are flagged and are forwarded to the editors. After verification, the new data are fed into the OdoBD database. The regions with poor data coverage are expected to be well documented with the help of enthusiastic citizen scientists.

The species data compiled with their respective eco-regions have a number of conservation applications. Conservation efforts can be undertaken at the regional scale using the eco-regions to distinguish distinct units of freshwater biodiversity. Our database may provide a crucial framework for identifying biogeographic locations that have the potential for being nominated as wetlands of international importance and are in need of protection. Similar processes to establish representative networks of protected freshwater areas by using eco-regions as a proxy, have been called for by IUCN World Conservation Congress, World Parks Congress and Convention on Biological Diversity (Abell *et al.* 2008). By compiling the list of less abundant, data deficient and near threatened species of Bangladesh our odonata database narrowed down the species required conservation attention. The main challenge remains the translation of these analyses into conservation implementation at local and national scales (da Fonseca *et al.* 2000). Conducting workshops providing local participants with biodiversity data to set up a consensus on individual conservation priorities is one of the most promising strategies for addressing the issue (Mittermeier *et al.* 1995).

In our study, we used the well-studied order of insects, Odonata, to construct a database of extant species in Bangladesh with their physical, ecological and genetic information. This database presents a useful source of information in determining the current state of Odonata communities in the region along with changes in species distribution. One of the major applications of this database is that it can be utilized as a data-exploration tool for comparative analysis. Comparison of odonata data from different regions may provide strong indications for global changes in biology as well as the underlying biological mechanisms for different habitat preferences of the odonates.

## Acknowledgements

We thank Ken Cheng for his comment on the earlier version of the manuscript. The digitization of the Odonata collection is supported by The Rufford Foundation, UK [grant number 18697-1] and The Explorers Club, USA. We also thank all the people, volunteers and citizen scientists for their help in the field-work, observations, sample collection and logistic support. We specially thank Borhan Uddin for helping us in the digitization of our dataset.

## Declarations

### Ethics approval and consent to participate

Not applicable

### Financial interest or benefit

Not applicable

